# miRNA analysis with *Prost!* reveals evolutionary conservation of organ-enriched expression and post-transcriptional modifications in three-spined stickleback and zebrafish

**DOI:** 10.1101/423533

**Authors:** Thomas Desvignes, Peter Batzel, Jason Sydes, B. Frank Eames, John Postlethwait

## Abstract

MicroRNAs (miRNAs) can have tissue-specific expression and functions; they can originate from dedicated miRNA genes, from non-canonical miRNA genes, or from mirror-miRNA genes and can also experience post-transcriptional variations. It remains unclear, however, which mechanisms of miRNA production or modification are tissue-specific and the extent of their evolutionary conservation. To address these issues, we developed the software *Prost!* (PRocessing Of Short Transcripts), which, among other features, allows accurate quantification of mature miRNAs, takes into account post-transcriptional processing, such as nucleotide editing, and helps identify mirror-miRNAs. Here, we applied *Prost!* to annotate and analyze miRNAs in three-spined stickleback (*Gasterosteus aculeatus*), a model fish for evolutionary biology reported to have a miRNome larger than most teleost fish. Zebrafish (*Danio rerio*), a distantly related teleost with a well-known miRNome, served as comparator. Despite reports suggesting that stickleback had a large miRNome, results showed that stickleback has 277 evolutionary-conserved *mir* genes and 366 unique mature miRNAs (excluding *mir430* gene replicates and the vaultRNA-derived *mir733*), similar to zebrafish. In addition, small RNA sequencing data from brain, heart, testis, and ovary in both stickleback and zebrafish identified suites of mature miRNAs that display organ-specific enrichment, which is, for many miRNAs, evolutionarily-conserved. These data also supported the hypothesis that evolutionarily-conserved, organ-specific mechanisms regulate miRNA post-transcriptional variations. In both stickleback and zebrafish, miR2188-5p was edited frequently with similar nucleotide editing patterns in the seed sequence in various tissues, and the editing rate was organ-specific with higher editing in the brain. In summary, *Prost!* is a critical new tool to identify and understand small RNAs and can help clarify a species’ miRNA biology, as shown here for an important fish model for the evolution of developmental mechanisms, and can provide insight into organ-specific expression and evolutionary-conserved miRNA post-transcriptional mechanisms.

## Introduction

microRNAs (miRNAs) are small non-coding RNA molecules about 20-22 nucleotides long that control gene expression post-transcriptionally by repressing translation or inducing the decay of targeted messenger RNA transcripts (mRNAs) ^1–3^. miRNAs participate in virtually all biological processes, including the control of cell specification, cell differentiation, organ development, and organ physiology ^4–6^ as well as pathologies in humans and other animals ^7–10^. miRNA genes also appear to be evolutionarily-conserved in number, sequence, and genomic environment across metazoans ^3,11–15^, but the evolutionary conservation of miRNA-specific functions remains incompletely understood.

Canonically, miRNA genes are transcribed into a primary transcript (pri-miRNA) that folds into a hairpin, from which the enzyme Drosha cleaves off the free 5′ and 3′ ends, thereby producing the precursor miRNA (pre-miRNA). The pre-miRNA, which assumes a stem-loop hairpin conformation, is then exported into the cytoplasm where a second enzyme, Dicer, trims off the loop and releases a miRNA duplex ^16,17^. One strand of the miRNA duplex is usually degraded, while the other strand loads into the RNA-Induced Silencing Complex (RISC), the effector of the miRNA regulation system. Once incorporated into the RISC, the miRNA drives the association of the enzymatic complex to specific mRNA transcripts by base pairing of the miRNA seed (nucleotides 2-8 from the 5′ end) to the targeted transcript’s 3′ UTR. Depending on pairing strength, the association of the RISC to the messenger RNA will either induce the decay of the transcript or prevent its translation. Other pathways and other gene types can also produce miRNAs (e.g. Drosha- or Dicer-independent pathways, introns of protein-coding genes (miRtrons), lncRNAs, snoRNAs ^18–22^).

Besides originating from a variety of biogenesis pathways and gene types, miRNA sequence variations can arise post-transcriptionally, resulting in variations in size and nucleotide sequence; these variants are called isomiRs ^2,23,24^. The most frequent post-transcriptional modification involves variations in length at the 3′ end of miRNAs. Length modifications at the 5′ end of miRNAs may occur less frequently because they cause a shift in the seed, which can modify the identity of targeted transcripts and, therefore, drastically change the miRNA’s function ^25^. miRNA sequence variation can also occur due to post-transcriptional editing, in which ADAR enzymes post-transcriptionally modify a nucleotide, usually an adenosine (A), into another base, usually an inosine (I) ^26–28^. These post-transcriptional modifications have now been shown to be physiologically relevant ^28–32^, but whether post-transcriptional editing occurs in a directed and regulated, organ-specific manner is still currently unknown.

The diversity of miRNAs, their variations, and the rapid expansion of small RNA sequencing reveal the need for small RNA analysis tools that can encompass the full diversity of gene origins and variations in miRNA sequences and extract information related to miRNA gene annotation in novel species, gene expression levels, strand selection from miRNA duplexes after Dicer processing, and potentially physiologically important post-transcriptional modifications.

Several bioinformatics tools are currently available to study miRNAs using small RNA sequencing datasets ^33–39^. Many tools start by filtering reads that can readily be annotated as miRNAs and then study their expression, sometimes using a genomic reference. Other tools have specialized in the discovery of novel miRNAs or the study of isomiRs. These tools often perform well for their respective functions, but in many cases, lack transparency in their filtering and annotating algorithms, have few user-defined parameter choices that might help tune a user’s specific application, and/or lack the ability to inspect the entire small RNA dataset and omit sequences not already annotated as a miRNA. With increasing amounts of data and read diversity, a more global approach was required to address sequencing output by analyzing every single read, even if they are not yet annotated as a type of coding or non-coding RNA. In addition, analysis tools should give attention to read alignments on a genomic reference to differentiate fragments potentially originating from one or multiple loci. While many tools are available to study small RNA sequencing datasets, current tools usually do not provide a comprehensive, genome-based, analysis of small RNA datasets, thus limiting the study of the full complexity of an experiment by missing out on some of the post-transcriptional processes affecting the diversity of small RNAs.

To help study the complexity of miRNA sequences in small RNA-seq data, we generated a new software tool *Prost!* (PRocessing Of Small Transcripts), which facilitates the identification of miRNAs for annotation, quantifies annotated miRNAs, and details variations (isomiRs) observed in each sample. *Prost!* is open-source software and is publicly available ^40^. Earlier versions of *Prost!* have been used to annotate zebrafish and spotted gar miRNAs ^41,42^, as well as to identify erythromiRs in white-blooded Antarctic icefish ^43^.

To investigate the evolutionary conservation of miRNAs in teleost fish, we performed small RNA sequencing on four organs (brain, heart, testis, and ovary) in two distantly related teleost laboratory model fishes: the zebrafish *Danio rerio* and the three-spined stickleback *Gasterosteus aculeatus*. While zebrafish miRNAs are well annotated ^41,44,45^, stickleback miRNAs aren’t, and current predicted annotations provide miRNA gene number estimations ranging from several hundred to well over a thousand genes ^46–49^, which is more than four times the number of miRNA genes in zebrafish. In addition, no study has so far investigated the potential conservation of miRNA functions across teleost fish species, or studied post-transcriptional modifications in teleost mature miRNAs. Here we thus addressed the following questions: 1) Is the evolutionarily-conserved stickleback miRNome significantly larger than that in other teleost species as reported? 2) Is organ-enrichment of miRNA expression shared by zebrafish and stickleback? And 3) Are post-transcriptional modifications controlled to display organ-specificity and are they shared by zebrafish and stickleback?

## Materials and methods

### Origin of sampled fish

Four reproductive adult zebrafish (*Danio rerio*, AB strain, two males and two females) were obtained from the University of Oregon Aquatic Animal Core Facility and four reproductive adult three-spined stickleback (*Gasterosteus aculeatus*, two males and two females) of a fresh water laboratory strain derived from Boot Lake, Alaska were obtained from Mark Currey in the W. Cresko Laboratory (University of Oregon). Animals were handled in accordance with good animal practice as approved by the University of Oregon Institutional Animal Care and Use Committee (Animal Welfare Assurance Number A-3009-01, IACUC protocol 12-02RA).

### RNA extraction and small RNA Library preparation

Immediately following euthanasia by overdose of MS-222, fin clips, brains, heart ventricles, and testes were sampled from two male zebrafish and two male stickleback, and fin clips and ovaries from two female zebrafish and two female stickleback. DNA was extracted from fin clips using a generic proteinase K DNA extraction method, and both small and large RNAs from each individual organ were extracted using Norgen Biotek microRNA purification kit according to the manufacturer’s instructions. Using the small RNA extract fractions, for each male of each species, we prepared three individual libraries (brain, heart ventricle, and testis), and for each female of each species we prepared a single library (ovary). In total, 16 small RNA libraries were then prepared and barcoded using the BiooScientific NEXTflex™ small RNA sequencing v1 kit with 15 PCR cycles. Libraries were sequenced on Illumina HiSeq2500 platform at the University of Oregon Genomics and Cell Characterization Core Facility (GC3F). Raw single-end 50-nt long reads were deposited in the NCBI Short Read Archive under project accession numbers SRP157992 and SRP039502 for stickleback and zebrafish, respectively.

### *Prost!* workflow

Raw reads from all sixteen libraries were pre-processed identically. Reads that did not pass Illumina’s chastity filter were discarded. Adapter sequences were trimmed from raw reads using cutadapt ^50^ with parameters: --overlap 10 -a TGGAATTCTCGGGTGCCAAGG --minimum-length 1. Reads were then quality filtered using fastq_quality_filter of the FASTX-Toolkit (http://hannonlab.cshl.edu/fastx_toolkit/commandline.html) (with parameters: -Q33 -q 30-p 100). The remaining reads were converted from FASTQ format to FASTA format.

The reads were then processed using *Prost!*, which is available online at https://prost.readthedocs.io and https://github.com/uoregon-postlethwait/prost ^40^. Briefly, *Prost!* size-selects reads for lengths typical of miRNAs and tracks the number of reads matching any given sequence. We configured *Prost!* to select for reads 17 to 25 nucleotides in length. *Prost!* then aligns the unique set of sequences to a reference genome using bbmapskimmer.sh of the BBMap suite (https://sourceforge.net/projects/bbmap/) (with parameters: mdtag=t scoretag=f inserttag=f stoptag=f maxindel=0 slow=t outputunmapped=f idtag=f minid=0.50 ssao=f strictmaxindel=t usemodulo=f cigar=t sssr=0.25 trimreaddescriptions=t secondary=t ambiguous=all maxsites=4000000 k=7 usejni=f maxsites2=4000000 idfilter=0.50). We configured *Prost!* to use the publicly available genome assemblies for three-spined stickleback (BROAD S1) and zebrafish (GRCz10) ^49^. *Prost!* uses the genome alignments to group sequences by genomic location. We configured *Prost!* to retain only sequences with a minimum of five identical reads for the initial annotation path, and only sequences with a minimum of 30 reads for the differential expression analysis.

*Prost!* then annotates the genomic location groups of reads by aligning against the mature and hairpin sequences of known miRNAs using bbmap.sh of the BBMap suite, as well as performing a reverse alignment of known mature sequences against the unique set of reads (with parameters: mdtag=t scoretag=f inserttag=f stoptag=f maxindel=0 slow=t outputunmapped=f idtag=f minid=0.50 ssao=f strictmaxindel=t usemodulo=f cigar=t sssr=0.25 trimreaddescriptions=t secondary=t ambiguous=all maxsites=4000000 k=7 usejni=f maxsites2=4000000 idfilter=0.50 nodisk). We configured *Prost!* to use all chordate mature and hairpin sequences present in miRBase Release 21 ^44^, the extended zebrafish miRNA annotation ^41^, and the spotted gar miRNA annotation ^42^. Gene nomenclature follows recent conventions ^2^, including those for zebrafish ^51^.

### Differential expression analyses

From *Prost!* output, we used the non-normalized read counts of annotated miRNA reads to perform differential expression analysis between organs by pair-wise comparisons using the DESeq2 package ^52^. For isomiR reads that could be variants of two or more miRNAs with equal probability, we partitioned their read counts proportionally based on counts of the annotated miRNAs from which they can differ. In addition, when a miRNA was not detected in an organ, a read count of one was used instead of zero to facilitate the calculation of an adjusted p-value for that miRNA. We selected the “local” type trend line fitting model (FitType) and used a stringent maximum adjusted p-value of 1% to consider miRNAs as differentially expressed between two organs. Each pairwise comparison was subsequently verified for appropriate p-value distributions and compatibility with the negative binomial probability model used by DESeq2 (Supplementary File1 and 2 for stickleback and zebrafish, respectively). Heat maps were generated using the Broad Institute Morpheus webserver ^3^ (https://software.broadinstitute.org/morpheus/) using log2-transformed normalized counts from annotated miRNAs that displayed a minimum normalized average expression of 5 Reads-per-Million (RPM) across all samples. Hierarchical clustering on both rows and columns was performed using the “one minus Pearson’s correlation” model and the “average” linkage method.

### Tissue-Specificity Index

To evaluate the organ-enrichment of the expression, we calculated for each miRNA a tissue specificity index (TSI) which is analogous to the TSI ‘tau’ for mRNAs ^54^ and has been previously used for miRNAs ^55^. The TSI varies from 0 to 1, with TSI close to 0 corresponding to miRNAs expressed in many or most tissues at similar levels (i.e. ‘housekeeping’ miRNAs), and TSI close to 1, corresponding to miRNAs expressed in a specific tissue (i.e. tissue-specific miRNAs).

### PCR analyses

To confirm editing events, we designed PCR primers to amplify primary miRNAs (pri-miRNAs) both from genomic DNA and from large RNA extracts of each investigated individual. This process allows the verification of putatively edited bases, rules out single nucleotide polymorphisms (SNPs) with respect to the reference genome sequence, and tests whether the transcribed pri-miRNA contains the edited base. Supplemental Table 1 contains primer sequences. PCR reactions were performed as previously described ^56^, and the product of each reaction was cleaned using Diffinity RapidTip (Diffinity Genomics, USA) and sequenced by Genewiz (South Plainfield, NJ, USA). Relative frequency of each base at various positions in the miRNAs were displayed using the WebLogo3 webserver ^57^. Putative miRNA targets were predicted using miRAnda 3.3a ^58,59^ with default parameters and the 3′ UTR annotations present in Ensembl release 79 genome assemblies (BROAD S1 for stickleback, Zv9 for zebrafish). Zebrafish to stickleback gene orthology was called by Ensembl biomart.

**Table 1:** Stickleback miRNA annotation statistics.

|  |  |
| --- | --- |
| Number of miRNA genes | 277 |
| Number of *mir430* genes (predicted by Ensembl) | 139 |
| Number of vaultRNA genes producing vault-derived miRNAs (i.e. *mir733* genes) | 3 |
| Total number of miRNA-producing genes | 418 |
| Number of mature miRNAs | 484 |
| Number of unique miRNAs | 366 |
| Number of miRNA genes with both 5p and 3p strands annotated | 214 (77%) |
| Number of miRNA genes with only the 5p mature strand annotated | 34 (12%) |
| Number of miRNA genes with only the 3p mature strand annotated | 16 (6%) |
| Number of miRNA genes with no mature strand annotated | 13 (5%) |

## Results and Discussion

### *Prost!*, a tool for analyzing small RNA sequencing reads

*Prost!* (PRocessing Of Short Transcripts) differs in three main ways from the majority of other tools developed to investigate small RNA-seq data. First, *Prost!* aligns reads to a user-defined genomic dataset (e.g. a genome assembly, Figure 1). This initial alignment permits retention of all sequencing reads that match, perfectly or with few errors, the “genomic dataset”, even if these matches are not yet known coding or non-coding RNA fragments. As such, *Prost!* enables the study of not only miRNAs, but also other small RNAs, such as piRNAs, t-RNA fragments, or the degradome of other RNA biotypes (e.g. snoRNAs, Y_RNA, vault-RNA). Second, *Prost!* groups reads based on their potential genomic origin(s), on their seed sequence, and ultimately on their annotation (Figure 1). This step allows the regrouping of sequence variants that could originate from one locus or from multiple loci. Conversely, this step discriminates reads that could only originate from a limited number of paralogous loci, increasing the understanding of gene expression and locus-specific expression levels. Third, *Prost!* analyzes in depth the subset of reads that had been annotated based on the user-provided annotation dataset (e.g. miRNA or piRNA) and reports frequencies of individual sequence variations with respect to both the reference genome and the most expressed sequence that aligns perfectly to the genome from a genomic location group or annotation group (Figure 1). This step ultimately provides a comprehensive report on potential post-transcriptional modifications for each group of sequences. *Prost!* was written in Python and takes as input a list of sequencing sample files. *Prost!* can be configured with a simple and well annotated configuration file and optional command line flags, allowing the user to optimize *Prost!* for each specific dataset, experimental design, and experimental goals (e.g. annotation or quantification). The output can be retrieved either as an individual report per analysis step as tab-separated value files, or combined into a single Excel file with each step provided as an individual tab that contains indexes, similar to primary-foreign keys of relational databases, facilitating the navigation from tab-to-tab to retrace and understand the entire analysis process (Figure 1). The supplementary files 3 and 4 are the *Prost!* output files used for differential expression analysis for stickleback and zebrafish, respectively.

**Figure 1:**
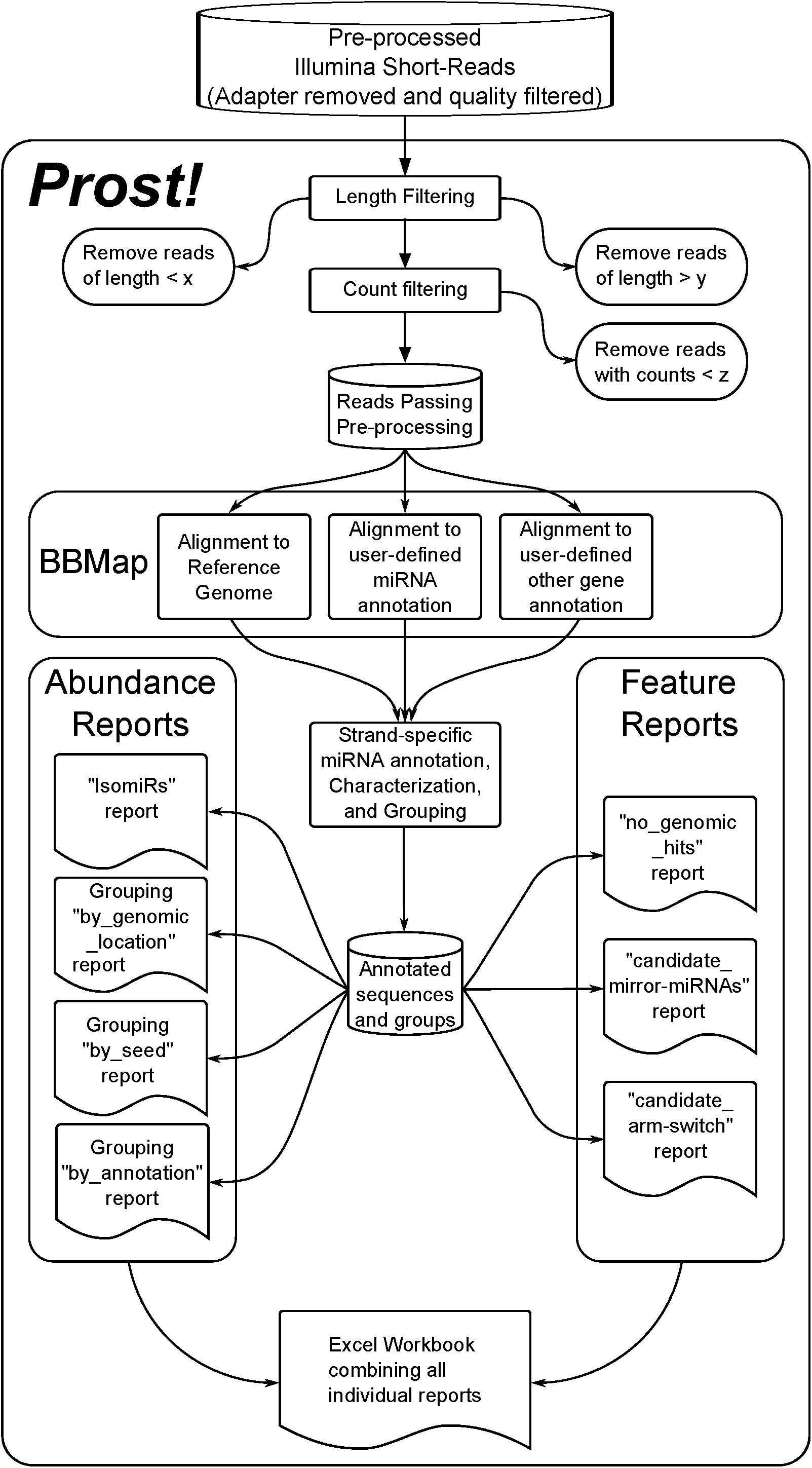
*Prost!* data processing flowchart. Flow chart displaying the input, pre-processing, categorization, alignment, and output report steps of *Prost!*

### Evolutionary conservation of the teleost miRNome in stickleback

ZooMir ^46^, Ensembl ^49^, and ^48^, predicted that stickleback has 483, 504, and 595 miRNA genes respectively, and ^47^ predicted 1486 mature miRNAs. Other well annotated teleost fish have substantially smaller miRNomes, consisting of about 250-350 genes ^44,45,49^. This discrepancy raises the question of whether the stickleback evolutionarily-conserved miRNome (miRNA genes that are conserved by at least two teleost species) is comparable to other well-annotated teleost genomes and contains approximately 250 to 300 miRNA genes, or whether the additional predicted stickleback miRNA genes are lineage-specific miRNAs and/or false predictions.

Using *Prost!* on our small RNA sequencing data of brain, heart, testis and ovary, we annotated 264 miRNA genes, with a total of 366 unique mature miRNAs (excluding the highly replicated *mir430* genes and the vaultRNA-derived *mir733*) (Table 1). Among these 264 miRNA genes, we were able to annotate both 5p and 3p strands for 214 genes (81%) and only one strand for 50 genes (19%) (Table 1). A search by sequence identity, orthology, and synteny conservation, for other known conserved teleost miRNAs ^41,44,45^ that were not among the 264 miRNA genes annotated with *Prost!* identified 13 more miRNA genes (Table 1). For these 13 miRNAs, however, because no mature miRNAs were present in our four-organ sequencing data, only the putative pri-miRNAs were annotated. Supplementary Table 2 combines all information of pri-miRNA and mature miRNA names, sequences, Ensembl Accession numbers if available, and position on the stickleback ‘BROAD S1’ genome assembly.

By comparing all available annotations, and excluding *mir430* genes and the vaultRNA-derived miRNAs *mir733* that form unique families of miRNA genes, we find that our stickleback annotation contained many of the miRNAs found in other stickleback annotations ^46,48,49^ and lacked some other genes. We did not include in our comparison the Chaturvedi et al ^47^ annotation because it was generated without strand-specificity. Compared to the other annotations (excluding 64, 139, and one *mir430* genes from ZooMir, Ensembl, and Rastorguev et al, respectively and one *mir733* genes from Rastorguev et al), 194 of 419 (46%) genes in ZoomiR, 234 of 593 genes (39%) in Rastorguev, and 250 of 365 genes (68%) in Ensembl overlapped with our annotation (Figure 2). Only three miRNAs were missing in our annotation that were present in at least two others, but all three of these (*mir204a, mir705*, and *mir1788*) are among the 13 known evolutionarily-conserved miRNAs that we annotated by orthology and for which no sequencing reads were present in our dataset, explaining why *Prost!* didn’t suggest an annotation for them. All the other miRNAs missing from our annotation were predicted in only one of the other three annotations (Figure 2), suggesting that predicted miRNAs our annotation lacks either correspond to false predictions, or are stickleback-specific genes, which *Prost!* wasn’t designed to detect.

**Figure 2:**
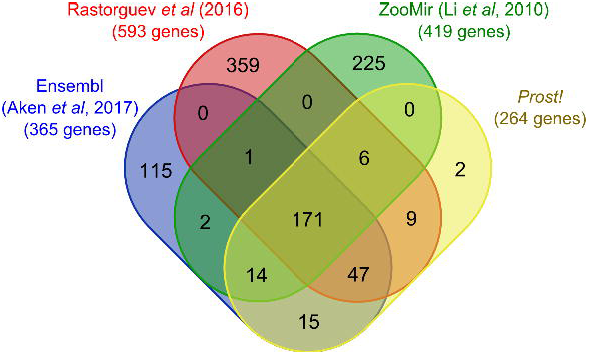
Overlap of the different existing stickleback miRNA annotations. Genomic locations of stickleback pre-miRNAs were retrieved from other stickleback annotations ^46,48,49^ and compared with each other and with those identified by *Prost!*. A pre-miRNA from one annotation was considered the same as a pre-miRNA from another annotation if they shared at least a 25 nucleotide overlap.

In summary, *Prost!* allowed us to efficiently annotate a complete set of evolutionarily-conserved miRNAs in stickleback and verify that the stickleback evolutionarily-conserved miRNome, with 277 miRNA genes, is similar to other well-annotated teleost species, which typically contains approximatively 250 to 300 miRNA genes (excluding the *mir430* genes). Supplementary files 5 and 6 provide annotation for the pri-miRNAs and unique mature miRNAs, respectively. The additional miRNAs previously predicted by others and missing from our annotation might be due to either over-identification in the other works or lineage-specific miRNAs, which our analysis using *Prost!* was not designed to identify.

### Identification of mirror-miRNAs in teleosts

In addition to the annotation of miRNA genes and mature miRNAs, *Prost!* facilitates rapid inspection of data to identify mirror-miRNA candidates by automatically filtering small RNA reads that originate from opposite DNA strands at the same location in the genome, so called mirror-miRNAs ^41,60,61^. Teleost fish have mirror-miRNAs and some are conserved across vertebrate species such as conserved mirror-miRNA pair *mir214*/*mir3120* in human and zebrafish, and the two other teleost mirror-miRNA pairs (*mir7547*/*mir7553* and *mir7552a*/*mir7552aos*) ^41^. In the list of candidate mirror-miRNAs generated by *Prost!*, we found the *mir214*/*mir3120* pair in both stickleback and zebrafish, while the two other known zebrafish mirror-miRNA pairs were not found in stickleback. Only miR7552a-5p, however, was present in our stickleback sequencing data, but given the limited number of organs studied, the mirror-miRNA pair mir7552a/mir7552aos might also be expressed in other stickleback organs. On the other hand, the pair *mir7547*/*mir7553* could not be found in the stickleback genome assembly by sequence homology or conserved synteny nor were they present in our sequencing data; we conclude that our data provides no evidence for this miRNA pair in stickleback. Nonetheless, *Prost!* allowed us to readily confirm the conservation of the mirror-miRNA pair *mir214*/*mir3120* in stickleback, demonstrating the genomic and transcriptomic conservation of these mirror-miRNAs among teleost fish.

### Organ-specific miRNA expression

miRNAs are generally considered to be specialized in function and to display organ- and even cell type-specific enrichment ^4,11,62^. Most of these data, however, are from mammals ^55,63–65^, so the extent to which this conservation of expression and function is similar among teleosts is unknown. We hypothesized that among stickleback and zebrafish miRNAs, a large proportion of the evolutionarily-conserved miRNAs should display organ-specific enrichment and tissue-specificity.

To show organ-specific enrichment of miRNAs in stickleback, we studied the expression of 258 mature miRNAs (of the 309 that *Prost!* detected) that displayed an average expression of at least 5 RPM across all eight stickleback samples. Pairwise differential expression analysis using the DESeq2 R package ^52^ showed that 1) the brain displayed the greatest number of differentially expressed (DE) miRNAs among the four studied organs, and 2) the gonads (testis and ovary), displayed the fewest DE miRNAs (Figure 3A).

**Figure 3:**
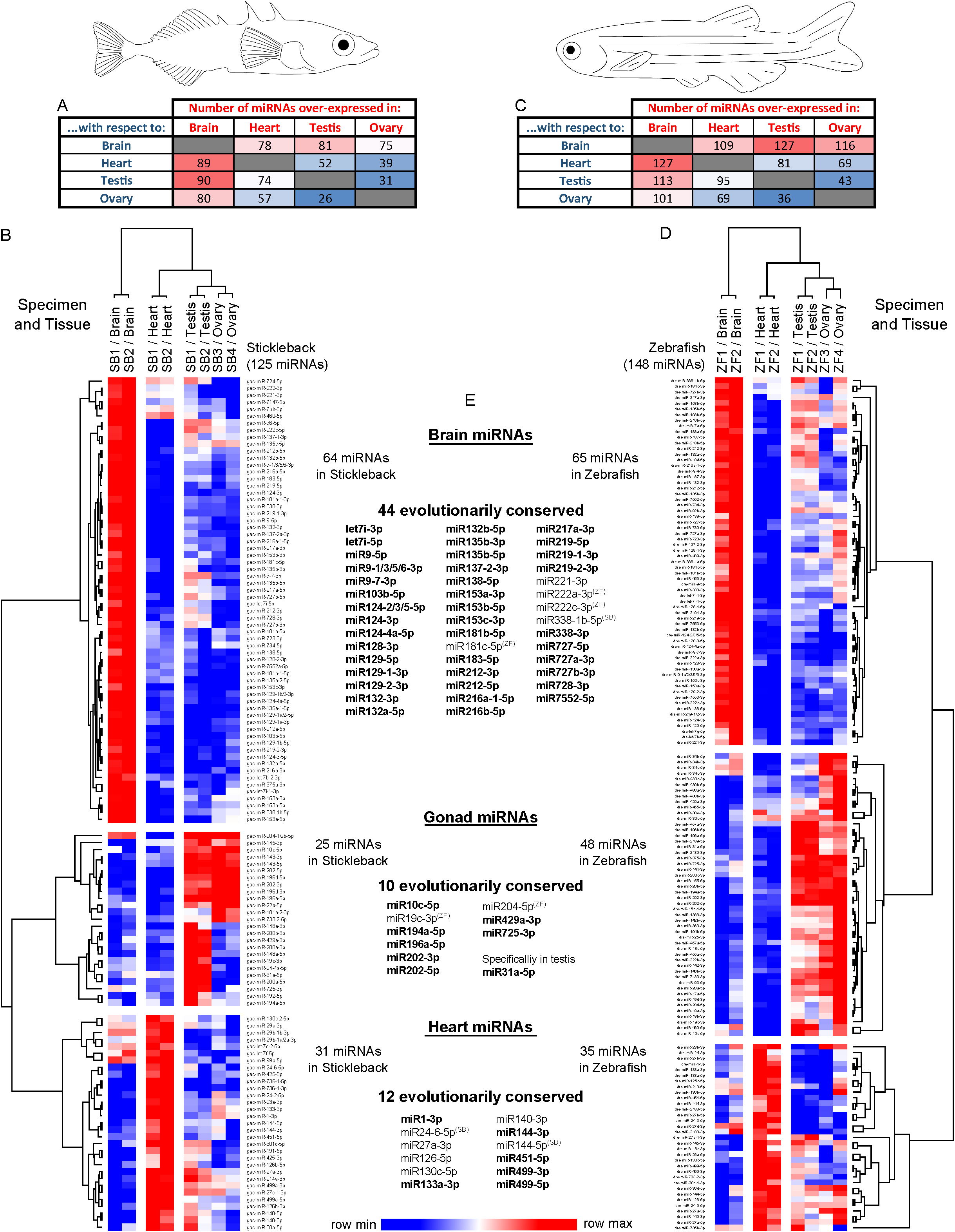
Differential expression and conservation of miRNAs in stickleback and zebrafish brain, heart, testis, and ovary. A) Heat map of the number of stickleback mature miRNAs over-expressed in each tissue compared to each other tissue. B) Heat map of the 125 stickleback mature miRNAs consistently enriched in one tissue compared to the three other tissues, or in gonads compared to brain and heart. C) Heat map of the number of zebrafish mature miRNAs over-expressed in each tissue compared to each other tissue. D) Heat map of the 148 zebrafish mature miRNAs consistently enriched in one tissue compared to the three other tissues, or in gonads compared to brain and heart. For all heat maps, blue indicates the lowest level of expression in the row and red indicates the highest level of expression in the row. E) Lists of tissue-enriched miRNAs that are evolutionarily-conserved between stickleback and zebrafish. Bold lettering denotes that the miRNA has a TSI > 0.85 in both species. Superscripted SB (Stickleback) or ZF (Zebrafish) denotes that this specific miRNA has a TSI > 0.85 in the corresponding species but not in the other.

In the six pairwise DE analyses, 64 miRNAs were consistently over-expressed in the stickleback brain compared to each of the three other organs, compared to only 31, 11 and eight for heart, testis, and ovary, respectively (Figure 3B, Supplementary Table 3). Because testis and ovary showed few organ-enriched DE miRNAs and share some common developmental processes in gametogenesis (e.g. meiosis), we looked at miRNAs that were over-expressed in both testis and ovary compared to both heart and brain. Six additional miRNAs were similarly enriched in both testis and ovaries compared to the other organs, bringing the total number of miRNAs that are enriched in one or both gonads to 25 (Supplementary Table 3). Altogether, 125 miRNAs (i.e. 48% of the 258 minimally expressed miRNAs) displayed organ enrichment in either brain, heart, testis, ovary or in both gonads, validating our hypothesis that many miRNAs in stickleback display organ-specific enrichment. More organ-enriched miRNAs would likely be identified with increasing number of organs studied.

To confirm this organ-enrichment, we analyzed the tissue specificity of each minimally expressed miRNA using the tissue specificity index (TSI), combining the testis and ovary data into a common ‘gonad’ tissue type. Of the 258 studied miRNAs, most (150; 58%) displayed intermediate specificity, implying that they were predominantly expressed in one or more organs but that expression in at least one other organ, was still significant (Figure 4A). 93 miRNAs (36%) showed a TSI > 0.85, which is considered a threshold for tissue-specificity ^54,55^, and only 15 miRNAs (6%) showed ubiquitous, similar expression levels among the studied tissues (Figure 4A). In addition, TSI scores agreed with the differential expression analysis, showing that the tissue-enriched miRNAs also tend to have the highest TSI scores (Figure 3B, Figure 4A). Among the DE miRNAs that also have a TSI > 0.85, some displayed clear enrichment in brain (e.g. miR9-5p, miR124-3p, and miR138-5p, Figure 4C-E), in testis (i.e. miR31a-3p, Figure 4F), in gonads (e.g. miR196a-5p and miR202-5p, Figure 4G-H), or in heart (e.g. miR1-3p, miR133-3p, miR499-5p, Figure 4I-K). Because we studied only four organs in this study, some miRNAs that we categorized as non-specific or enriched in only one tissue might be enriched in other organs. For example, miR122-5p, which is known to be mostly expressed in liver in vertebrates ^45,55^, showed low, non-specific expression in all four organs we investigated with an average of 18 RPM and a TSI score of 0.53.

**Figure 4:**
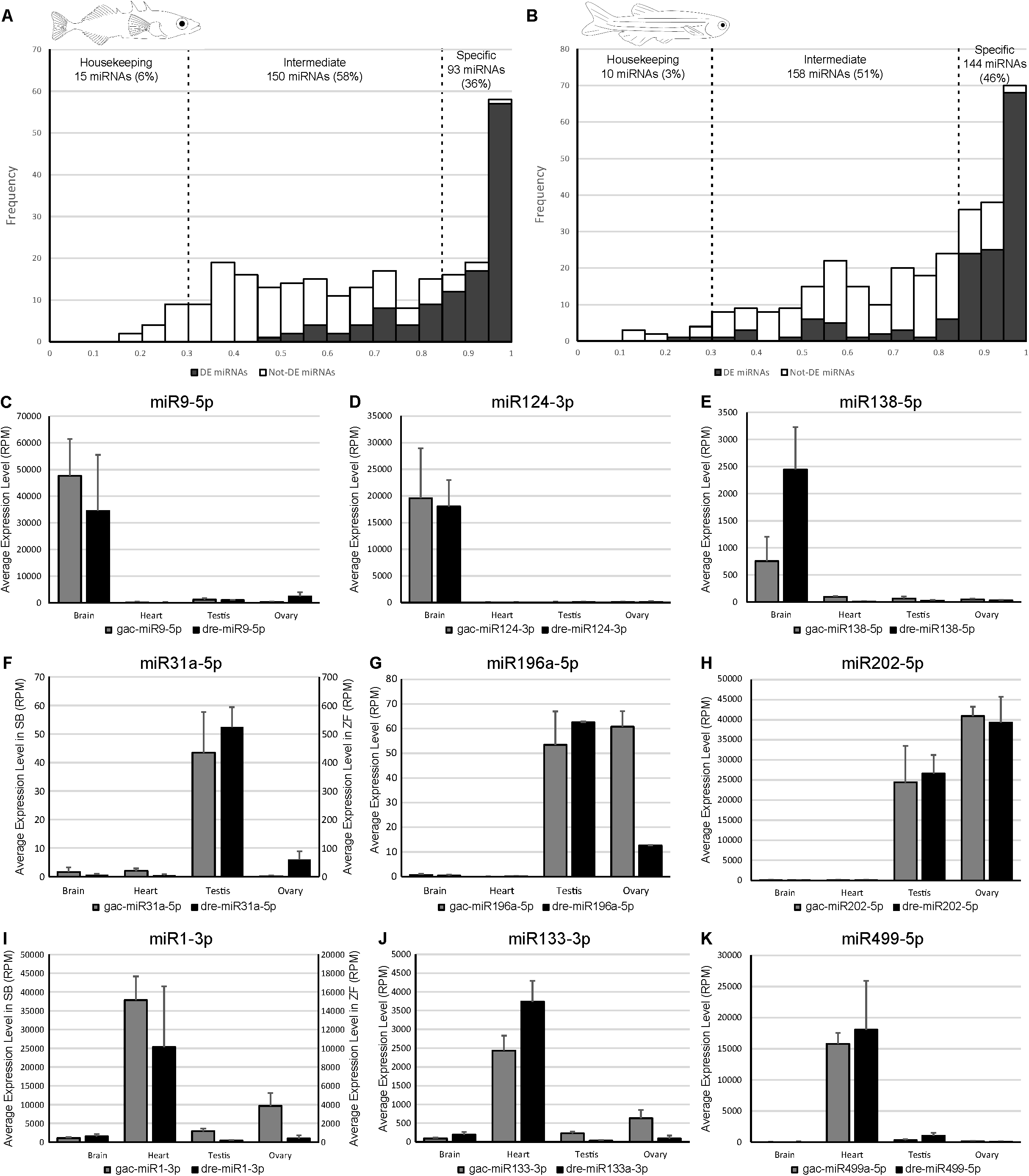
miRNA organ-specific expression. A-B) Frequency plot of TSI values for all minimally-expressed stickleback and zebrafish miRNAs. Grey bars represent miRNAs that were also found to be enriched in brain, heart, testis, ovary, or in both gonads, and white bars represent miRNAs that were not found to be enriched in a specific tissue. C-K) Average tissue expression of evolutionarily-conserved, tissue-enriched miRNAs that have a TSI > 0.85. Expression levels are given in RPM (Reads per Million) with associated standard deviations for the four organs studied in both stickleback (grey bars) and zebrafish (black bars).

To identify organ-specific enrichment of miRNAs in zebrafish, we studied the expression of 312 mature miRNAs (of the 398 that *Prost!* detected) that displayed an average expression of at least 5 RPM across all eight zebrafish samples. Similar to stickleback, the brain had the most differently expressed miRNAs, and ovary and testis had the least (Figure 3C).

In all zebrafish pairwise comparisons, 65 miRNAs were consistently enriched in brain, 35 in heart, eight in testis, 22 in ovary, and an additional 18 miRNAs equally enriched in both gonads (Figure 3D, Supplementary Table 4). Altogether, 148 miRNAs (47% of the 312 minimally expressed zebrafish miRNAs) displayed organ-enrichment in brain, heart, testis, ovary, or in both gonads in zebrafish. Similar to the stickleback TSI analysis, of the 312 zebrafish miRNAs studied, most miRNAs (158; 51%) displayed intermediate specificity, 10 miRNAs (3%) showed overall ubiquitous expression levels among the studied tissues, and 144 miRNAs (46%) showed tissue-specificity (TSI > 0.85) (Figure 4A). Also, similar to stickleback, we observed that miRNAs identified as organ-enriched by differential expression analyses are among the miRNAs that have the highest TSI (Figure 3D, Figure 4B).

Taken together, results demonstrated that a large proportion of miRNAs displayed enrichment in a single organ in both stickleback and zebrafish (48% and 47%, respectively) and tissue-specific TSI scores (36 and 46% in stickleback and zebrafish, respectively). These results confirm that teleost miRNAs display a high level of tissue specificity, consistent with other vertebrates.

### Organ-enriched miRNAs are conserved between stickleback and zebrafish

The hypothesis that miRNA functions are conserved predicts that at least some of the organ-enriched miRNAs in stickleback would also be enriched in the same organ in zebrafish. To test this prediction, we compared the list of organ-enriched miRNAs in stickleback and zebrafish and found that 44 miRNAs were brain-enriched in both species (Figure 3B, D, E), with many of them already known to be brain-associated miRNAs in several fish species ^66–68^, for example miR9-5p, miR124-3p, and miR138-5p (Figure 4C-E), which are also highly expressed in brain and nervous organs in mammals ^45,55,69–73^. These observations suggest a strong evolutionary conservation of function of brain-related, miRNAs among vertebrates. Heart also displayed a significant number of evolutionarily-conserved, organ-enriched miRNAs (12 miRNAs) (Figure 3B, D, E). Heart-enriched miRNAs included mature products of the well-described vertebrate cardiac myomiR genes *mir1, mir133*, and *mir499* ^55,67,68,74–77^ and erythromiRs *mir144* and *mir451* ^43^. The former group participate in muscle formation and function, and the latter may reflect the presence of red blood cells in the heart ventricle at the time of RNA extraction.

Surprisingly, gonad-specific miRNAs appeared to be less conserved. Only miR31a-5p was found to be testis-enriched in both stickleback and zebrafish (Figure 3B, D, E, Figure 4F), while no miRNAs were ovary-enriched in both species (Figure 3B, D, E). In chicken, *Mir31* has been hypothesized to be involved in gonadal sex differentiation, because it is significantly more expressed in testes compared to ovaries at early sexual differentiation stages ^78^. In human, *MIR31* is down-regulated in the testis of an infertile adult human patient ^79^. While *mir31* has never been associated with either gonad differentiation or testicular function in fish, our data suggest a conserved role of *mir31* in testicular function among various vertebrate lineages. In addition, nine shared miRNAs were enriched in one or both gonads in both species (Figure 3B, D, E), potentially reflecting a role in reproduction in one or both sexes in both species. Interestingly, among the gonad-associated miRNAs in stickleback and zebrafish, most displayed species-specific tissue enrichment. For example, miR429a-3p was enriched in testis in stickleback but enriched in ovary in zebrafish; miR10c-5p was enriched in ovary in stickleback but enriched in testis in zebrafish; miR204-5p was enriched in ovary in zebrafish but enriched in both gonads in stickleback; miR196a-5p was enriched in testis in zebrafish and in both gonads in stickleback (Figure 4G); miR19c-3p, miR194a-5p, and miR725-3p were enriched in testis in stickleback but enriched in both gonads in zebrafish. In the case of the well-known gonad-specific miR202-5p ^80–82^, the expression level in the stickleback ovary was significantly higher than in testis; although the trend was the same in zebrafish, the difference was not statistically significant (Figure 4H).

The relatively weak evolutionary conservation of sex-specific gonad enrichment in teleost fish is surprising and suggests reduced selective pressure on their function compared to other organ-enriched miRNAs, and/or that differences in the control of reproduction exist between zebrafish and stickleback. Not enough information is currently available to distinguish between these two non-exclusive hypotheses. The large range of variations in sex-determination mechanisms, reproductive systems, reproductive state, and frequency of reproduction in teleost fish ^83,84^, however, might help explain the weak conservation of stickleback and zebrafish miRNA expression in gonads. Indeed, the miRNA regulation system might be evolving with each species’ reproductive biology and its associated genetic regulation. Some ancestral functions of a miRNA could be conserved in one species, could have evolved novel targets and regulatory pathways in another, or may simply be lost in a lineage-specific fashion. For example, a gonad-enriched ancestral miRNA might have specialized in testis in one lineage, while remaining gonad-enriched or becoming ovary-enriched in another lineage, as could have happened with miR10c-5p or miR429a-5p in our data. The study of more teleost species and inclusion of an outgroup to represent the common ancestor, such as spotted gar, is necessary to test these hypotheses.

### Evolutionarily-conserved brain-specific post-transcriptional miRNA seed editing

Post-transcriptional modification of miRNAs is frequent and generally originates from variation in biogenesis or enzyme-catalyzed nucleotide modification. These modifications result in groups of related sequences, isomiRs, which are different mature miRNA sequences originating from the same gene ^2,23,24^.

To study post-transcriptional modifications in teleost fish and to ask whether they are evolutionarily-conserved, we developed a feature in *Prost*! to calculate and color-code each type of post-transcriptional modification at individual genomic loci. Results showed that in stickleback, miR2188-5p displayed a high frequency of seed-editing in the brain (34%), compared to other organs (9.6%) (Figure 5A). In zebrafish, similar analyses of post-transcriptional modifications revealed that the same miRNA, miR2188-5p, also displayed a higher rate of seed editing in the brain (12%), compared to other tissues (1.0%; Figure 5A). *Prost!* output also revealed that the stickleback miR2188-5p isomiR pool was composed of sequences displaying three different seeds: the genome-encoded seed (67% of the sequences), a seed with an adenosine-to-guanosine (A-to-G) substitution at position 8 of the miRNA (nt8; 26% of the sequences), and a seed with two A-to-G substitutions, one at nucleotide 2 and one at nucleotide 8 (nt2-nt8; 7% of the sequences) (Figure 5B). Examination of seed variations in zebrafish identified two different seeds: the genome-encoded seed (89% of the sequences) and a seed with an A-to-G substitution at the second nucleotide of the mature miRNA (nt2; 11% of the sequences) (Figure 5B). Because inosine bases are replaced by guanosine bases during the cDNA synthesis step of the sequencing library preparation protocols, sequencers report a guanosine where an inosine could have originally been present in the RNA molecule. Therefore, the nucleotides sequenced as guanosine in place of an adenosine in our sequencing data were likely to be post-transcriptionally ADAR-edited Inosine nucleotides.

**Figure 5:**
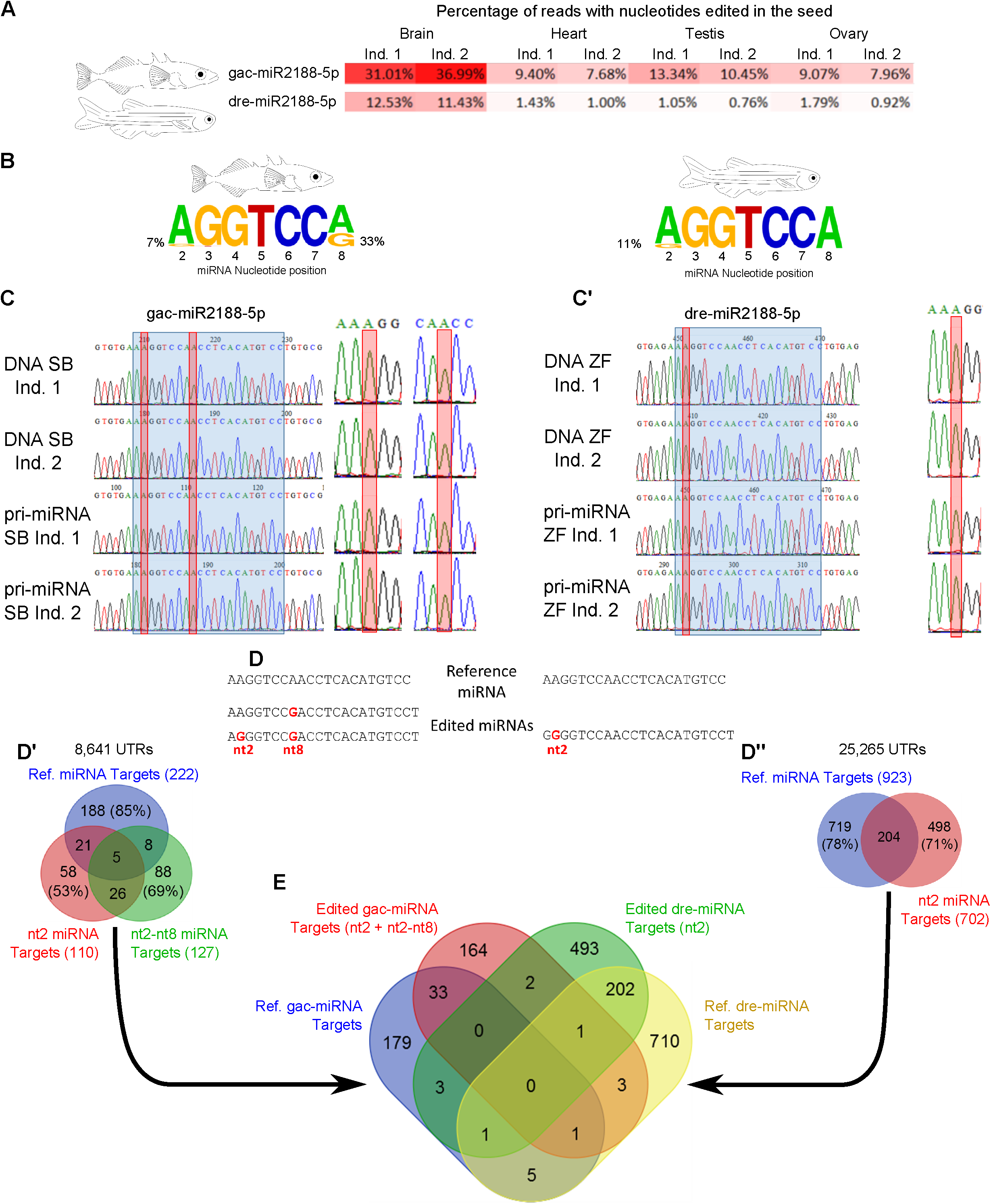
Evolutionarily-conserved brain-specific seed-editing of miR2188-5p. A) miR2188-5p seed editing frequency is higher in brain compared to other organs in both stickleback and zebrafish. B) Frequency of seed variants generated using WebLogo3 webserver. C) Sanger sequencing of DNA and pri-miRNAs of both stickleback specimens. C’) Sanger sequencing of DNA and pri-miRNAs of both zebrafish specimens. The blue box highlights the mature miRNA and the red box highlights the editing site. D) Mature miR2188-5p sequences used for target prediction in both stickleback and zebrafish. D’) Overlap of the predicted target mRNA sets for each mature miR2188-5p isomiR in stickleback. D’’) Overlap of the predicted target mRNA sets for each mature miR2188-5p isomiR in zebrafish. E) Overlap of predicted target mRNA sets for either genetically encoded or edited mature miR2188-5p in both stickleback and zebrafish.

To verify that the A-to-G substitutions are due to ADAR editing instead of potential miRNA allelic variations or other encoded SNPs, we sequenced the *mir2188* gene from genomic DNA of each stickleback and zebrafish individual that we had processed for small RNA sequencing. In both species, the genomes of both fish individuals used in our gene expression experiments contained the same nucleotides at the seed editing sites as the reference genome sequences (Figure 5C-C’). Therefore, the variations in sequence observed in the miR2188-5p miRNA seed were not genetically encoded, and thus likely resulted from post-transcriptional modifications. To confirm this result, because ADAR is enzymatically active only on RNA molecules that are double-stranded such as when miRNAs are in the precursor or duplex form ^26,28^, we sequenced pri-miRNAs from large RNA extracts of these same samples. In each case, we found that *pri-miR2188* was free of nucleotide substitutions at the second and/or eighth nucleotide of the mature miRNA (Figure 5C-C’). The post-transcriptional modifications altering the seed sequence in miR2188-5p were thus not genetically encoded, but occurred after the processing of the hairpin, possibly at the duplex stage. We conclude that post-transcriptional seed editing of miR2188-5p is organ-enriched and evolutionarily-conserved. This represents, to our knowledge, the first example of teleost organ-enriched evolutionarily-conserved seed editing.

miRNA seed editing, by changing the seed sequence, can alter the set of targeted transcripts and therefore modify the miRNA function ^28–32^. To evaluate the potential biological effects of miR2188-5p seed editing, we used miRAnda ^58,59^ and 3′ UTR sequences to predict mRNA targets of both the reference miRNA and the edited miRNAs (Figure 5D). For stickleback, the miR2188-5p isomiRs had few overlapping predicted target genes; in most cases, putative targets were unique to one of the three isomiRs (Figure 5D’). For zebrafish, the targets predicted for each isomiR were also largely non-overlapping, with less than 15% of predicted mRNA targets being in common for both isomiRs (Figure 5D’’).

Finally, an analysis of overlaps among putative mRNA target sets for stickleback and zebrafish miR2188-5p isomiRs showed that two transcripts were not predicted targets of the genomically encoded isomiR but were predicted targets of the seed-edited isomiRs in both stickleback and zebrafish. One was *cntnap3* (*contactin associated protein like 3*, ENSDARG00000067824), a cell adhesion molecule of unknown function in teleost fish. In human and mouse, however, *CNTNAP3* is known to be expressed in brain and spinal cord ^85,86^, and its dysregulation in developing mice impairs motor learning ^87^ and social behavior ^88^. In addition, high CNTNAP3 levels have been associated with schizophrenia in human ^89^. The second conserved target of the seed-edited miR2188-5p is *pdha1a* (*pyruvate dehydrogenase alpha 1*, ENSDARG00000012387), which is strongly expressed in the brain of developing zebrafish embryos ^90^. *PDHA1* mutations in human cause acid lactic buildup, resulting in impaired psychomotor development and chronic neurologic dysfunction with structural abnormalities in the central nervous system (OMIM 300502, ORPHA:79243). To our knowledge, no publications have identified miR2188-5p as an edited miRNA nor implicated it in brain function, but our data argue for more research. Study of multiple species is needed to understand the phylogenetic conservation of the brain-specific seed-editing of miR2188-5p and its potential role in vertebrate brain development and physiology.

## Conclusions

Results reported here show that the novel software *Prost!* facilitated the annotation of a set of evolutionary-conserved miRNAs in stickleback and verified that the stickleback miRNome, with 277 evolutionary-conserved miRNA genes, is comparable to other teleost species. In addition, as predicted, the differential expression analysis of miRNAs in four organs from adult stickleback and zebrafish revealed that about 48% and 47%, respectively, of minimally expressed miRNAs displayed significant organ-enrichment in either brain, heart, testis, ovary, or both gonads. Furthermore, supporting the hypothesis that organ-specific miRNAs are evolutionarily-conserved, enriched expression of specific miRNAs was found in both brain and heart of stickleback and zebrafish. In ovary and testis, however, fewer expressed miRNAs were conserved, although several miRNAs that were enriched in both gonads of one species tended to be enriched in at least one of the two gonad types in the other species. Finally, we demonstrated the conservation of organ-specific miR2188-5p seed editing in the brains of both zebrafish and stickleback, suggesting conservation of organ-specific, post-transcriptional miRNA modifications among teleosts.

## Authors’ contributions

Study concept and design: TD, BFE, PB, JS, JHP

Software development: PB, TD, JS, BFE, JHP

Acquisition of data: TD

Analysis and interpretation of data: TD, JS, PB

Wrote the manuscript: TD

Critical revision of the manuscript: TD, BFE, PB, JS, JHP

Obtained funding: JHP, TD

Study supervision: TD, JHP

## Acknowledgments

We thank the University of Oregon Aquatics Facility, Trevor Enright and Tim Mason for zebrafish care; Mark Currey and Bill Cresko for the gift of stickleback. We also thank Teretha Taylor (Albany State University, GA; UO SPUR Program 2R25HD070817) and Michael J. Beam who assisted in the early steps of stickleback miRNA annotation, and John Willis, David Clouthier, Kristin Artinger, Julien Bobe, Jérôme Montfort, Joanna Kelley, Igor Babiak, Leonardo M Martin, Teshome T Bizuayehu, Peter Alestrom, and Havard Aanes for constructive comments during the development of *Prost!*., as well as Brian Bushnell for helpful advice and insight about BBMap This work benefited from access to the University of Oregon high performance computers Talapas and ACISS (OCI-0960354).

## Supplementary Datasets

**Supplementary File 1:** Pairwise differential expression graphs in stickleback organs.

**Supplementary File 2:** Pairwise differential expression graphs in zebrafish organs.

**Supplementary File 3:** *Prost!* output file used for differential expression analysis for stickleback.

**Supplementary File 4:** *Prost!* output file used for differential expression analysis for stickleback.

**Supplementary File 5:** Stickleback primary miRNA annotation.

**Supplementary File 6:** Stickleback mature miRNA annotation.

**Supplementary Table 1:** PCR primers used for the study of miR2188-5p.

**Supplementary Table 2:** Complete stickleback miRNA annotation.

**Supplementary Table 3:** Tissue-specific lists of differentially expressed miRNAs in stickleback organs.

**Supplementary Table 4:** Tissue-specific lists of differentially expressed miRNAs in zebrafish organs.

## References

1. Carthew, R. W. & Sontheimer, E. J. Origins and Mechanisms of miRNAs and siRNAs. Cell 136, 642–655 (2009).

2. Desvignes, T. et al. miRNA Nomenclature: A View Incorporating Genetic Origins, Biosynthetic Pathways, and Sequence Variants. Trends Genet. 31, 613–626 (2015).

3. Bartel, D. P. Metazoan MicroRNAs. Cell 173, 20–51 (2018).

4. Christodoulou, F. et al. Ancient animal microRNAs and the evolution of tissue identity. Nature 463, 1084–1088 (2010).

5. Kosik, K. S. MicroRNAs and Cellular Phenotypy. Cell 143, 21–26 (2010).

6. Ameres, S. L. & Zamore, P. D. Diversifying microRNA sequence and function. Nat. Rev. Mol. Cell Biol. 14, 475–488 (2013).

7. Bhaskaran, M. & Mohan, M. MicroRNAs: History, Biogenesis, and Their Evolving Role in Animal Development and Disease. Vet. Pathol. 51, 759–774 (2014).

8. Leva, G. D., Garofalo, M. & Croce, C. M. MicroRNAs in Cancer. Annu. Rev. Pathol. Mech. Dis. 9, 287–314 (2014).

9. Ranganathan, K. & Sivasankar, V. MicroRNAs - Biology and clinical applications. J. Oral Maxillofac. Pathol. 18, 229 (2014).

10. Tüfekci, K. U., Öner, M. G., Meuwissen, R. L. J. & Genç, Ş. The Role of MicroRNAs in Human Diseases. in miRNomics: MicroRNA Biology and Computational Analysis 33–50 (Humana Press, Totowa, NJ, 2014).

11. Lee, C.-T., Risom, T. & Strauss, W. M. Evolutionary Conservation of MicroRNA Regulatory Circuits: An Examination of MicroRNA Gene Complexity and Conserved MicroRNA-Target Interactions through Metazoan Phylogeny. DNA Cell Biol. 26, 209–218 (2007).

12. Grimson, A. et al. Early origins and evolution of microRNAs and Piwi-interacting RNAs in animals. Nature 455, 1193–1197 (2008).

13. Wheeler, B. M. et al. The deep evolution of metazoan microRNAs. Evol. Dev. 11, 50–68 (2009).

14. Loh, Y.-H. E., Yi, S. V. & Streelman, J. T. Evolution of microRNAs and the diversification of species. Genome Biol. Evol. 3, 55–65 (2011).

15. Tarver, J. E., Donoghue, P. C. J. & Peterson, K. J. Do miRNAs have a deep evolutionary history? BioEssays 34, 857–866 (2012).

16. Winter, J., Jung, S., Keller, S., Gregory, R. I. & Diederichs, S. Many roads to maturity: microRNA biogenesis pathways and their regulation. Nat. Cell Biol. 11, 228–234 (2009).

17. Ha, M. & Kim, V. N. Regulation of microRNA biogenesis. Nat. Rev. Mol. Cell Biol. 15, 509–524 (2014).

18. Berezikov, E., Chung, W.-J., Willis, J., Cuppen, E. & Lai, E. C. Mammalian Mirtron Genes. Mol. Cell 28, 328–336 (2007).

19. Ruby, J. G., Jan, C. H. & Bartel, D. P. Intronic microRNA precursors that bypass Drosha processing. Nature 448, 83–86 (2007).

20. Cheloufi, S., Dos Santos, C. O., Chong, M. M. W. & Hannon, G. J. A dicer-independent miRNA biogenesis pathway that requires Ago catalysis. Nature 465, 584–589 (2010).

21. Cifuentes, D. et al. A Novel miRNA Processing Pathway Independent of Dicer Requires Argonaute2 Catalytic Activity. Science 328, 1694–1698 (2010).

22. Scott, M. S. & Ono, M. From snoRNA to miRNA: Dual function regulatory non-coding RNAs. Biochimie 93, 1987–1992 (2011).

23. Neilsen, C. T., Goodall, G. J. & Bracken, C. P. IsomiRs – the overlooked repertoire in the dynamic microRNAome. Trends Genet. 28, 544–549 (2012).

24. Guo, L. & Chen, F. A challenge for miRNA: multiple isomiRs in miRNAomics. Gene 544, 1–7 (2014).

25. Tan, G. C. et al. 5IZ isomiR variation is of functional and evolutionary importance. Nucleic Acids Res. 42, 9424–9435 (2014).

26. Yang, W. et al. Modulation of microRNA processing and expression through RNA editing by ADAR deaminases. Nat. Struct. Mol. Biol. 13, 13–21 (2006).

27. Heale, B. S. E. et al. Editing independent effects of ADARs on the miRNA/siRNA pathways. EMBO J. 28, 3145–3156 (2009).

28. Nishikura, K. A-to-I editing of coding and non-coding RNAs by ADARs. Nat. Rev. Mol. Cell Biol. 17, 83–96 (2016).

29. Kawahara, Y., Zinshteyn, B., Chendrimada, T. P., Shiekhattar, R. & Nishikura, K. RNA editing of the microRNA-151 precursor blocks cleavage by the Dicer–TRBP complex. EMBO Rep. 8, 763–769 (2007).

30. Kawahara, Y. et al. Redirection of Silencing Targets by Adenosine-to-Inosine Editing of miRNAs. Science 315, 1137–1140 (2007).

31. Kume, H., Hino, K., Galipon, J. & Ui-Tei, K. A-to-I editing in the miRNA seed region regulates target mRNA selection and silencing efficiency. Nucleic Acids Res. gku662 (2014). doi:10.1093/nar/gku662

32. Warnefors, M., Liechti, A., Halbert, J., Valloton, D. & Kaessmann, H. Conserved microRNA editing in mammalian evolution, development and disease. Genome Biol. 15, R83 (2014).

33. Pantano, L., Estivill, X. & Martí, E. SeqBuster, a bioinformatic tool for the processing and analysis of small RNAs datasets, reveals ubiquitous miRNA modifications in human embryonic cells. Nucleic Acids Res. 38, e34–e34 (2010).

34. Friedländer, M. R., Mackowiak, S. D., Li, N., Chen, W. & Rajewsky, N. miRDeep2 accurately identifies known and hundreds of novel microRNA genes in seven animal clades. Nucleic Acids Res. 40, 37–52 (2012).

35. Barturen, G. et al. sRNAbench: profiling of small RNAs and its sequence variants in single or multi-species high-throughput experiments. Methods Gener. Seq. -1, (2014).

36. Baras, A. S. et al. miRge-A Multiplexed Method of Processing Small RNA-Seq Data to Determine MicroRNA Entropy. Plos ONE 10, e0143066 (2015).

37. Rueda, A. et al. sRNAtoolbox: an integrated collection of small RNA research tools. Nucleic Acids Res. 43, W467–W473 (2015).

38. Lukasik, A., Wójcikowski, M. & Zielenkiewicz, P. Tools4miRs – one place to gather all the tools for miRNA analysis. Bioinformatics 32, 2722–2724 (2016).

39. Urgese, G., Paciello, G., Acquaviva, A. & Ficarra, E. isomiR-SEA: an RNA-Seq analysis tool for miRNAs/isomiRs expression level profiling and miRNA-mRNA interaction sites evaluation. BMC Bioinformatics 17, 148 (2016).

40. Batzel, P., Desvignes, T., Postlethwait, J. H., Eames, B. F. & Sydes, J. Prost!, a tool for miRNA annotation and next generation smallRNA sequencing experiment analysis. Zenodo (2015). doi:10.5281/zenodo.35422

41. Desvignes, T., Beam, M. J., Batzel, P., Sydes, J. & Postlethwait, J. H. Expanding the annotation of zebrafish microRNAs based on small RNA sequencing. Gene 546, 386–389 (2014).

42. Braasch, I. et al. The spotted gar genome illuminates vertebrate evolution and facilitates human-teleost comparisons. Nat. Genet. 48, 427–437 (2016).

43. Desvignes, T., Detrich III, H. W. & Postlethwait, J. H. Genomic conservation of erythropoietic microRNAs (erythromiRs) in white-blooded Antarctic icefish. Mar. Genomics 30, 27–34 (2016).

44. Kozomara, A. & Griffiths-Jones, S. miRBase: annotating high confidence microRNAs using deep sequencing data. Nucleic Acids Res. 42, D68–D73 (2013).

45. Fromm, B. et al. A Uniform System for the Annotation of Vertebrate microRNA Genes and the Evolution of the Human microRNAome. Annu. Rev. Genet. 49, 213–242 (2015).

46. Li, S.-C. et al. Identification of homologous microRNAs in 56 animal genomes. Genomics 96, 1–9 (2010).

47. Chaturvedi, A., Raeymaekers, J. A. M. & Volckaert, F. A. M. Computational identification of miRNAs, their targets and functions in three-spined stickleback (Gasterosteus aculeatus). Mol. Ecol. Resour. 14, 768–777 (2014).

48. Rastorguev, S. M. et al. Identification of novel microRNA genes in freshwater and marine ecotypes of the three-spined stickleback (Gasterosteus aculeatus). Mol. Ecol. Resour. 16, 1491–1498 (2016).

49. Aken, B. L. et al. Ensembl 2017. Nucleic Acids Res. 45, D635–D642 (2017).

50. Martin, M. Cutadapt removes adapter sequences from high-throughput sequencing reads. EMBnet.journal 17, 10–12 (2011).

51. Bradford, Y. et al. ZFIN: enhancements and updates to the zebrafish model organism database. Nucleic Acids Res. 39, D822–D829 (2011).

52. Love, M. I., Huber, W. & Anders, S. Moderated estimation of fold change and dispersion for RNA-seq data with DESeq2. Genome Biol. 15, 550 (2014).

53. Morpheus. Available at: https://software.broadinstitute.org/morpheus/.

54. Yanai, I. et al. Genome-wide midrange transcription profiles reveal expression level relationships in human tissue specification. Bioinformatics 21, 650–659 (2005).

55. Ludwig, N. et al. Distribution of miRNA expression across human tissues. Nucleic Acids Res. 44, 3865–3877 (2016).

56. Desvignes, T., Contreras, A. & Postlethwait, J. H. Evolution of the miR199-214 cluster and vertebrate skeletal development. RNA Biol. 11, 281–294 (2014).

57. Crooks, G. E., Hon, G., Chandonia, J.-M. & Brenner, S. E. WebLogo: A Sequence Logo Generator. Genome Res. 14, 1188–1190 (2004).

58. Enright, A. J. et al. MicroRNA targets in Drosophila. Genome Biol. 5, R1 (2003).

59. Betel, D., Wilson, M., Gabow, A., Marks, D. S. & Sander, C. The microRNA.org resource: targets and expression. Nucleic Acids Res. 36, D149–D153 (2008).

60. Tyler, D. M. et al. Functionally distinct regulatory RNAs generated by bidirectional transcription and processing of microRNA loci. Genes Dev. 22, 26–36 (2008).

61. Scott, H. et al. MiR-3120 Is a Mirror MicroRNA That Targets Heat Shock Cognate Protein 70 and Auxilin Messenger RNAs and Regulates Clathrin Vesicle Uncoating. J. Biol. Chem. 287, 14726–14733 (2012).

62. Halushka, M. K., Fromm, B., Peterson, K. J. & McCall, M. N. Big Strides in Cellular MicroRNA Expression. Trends Genet. 34, 165–167 (2018).

63. De Rie, D. et al. An integrated expression atlas of miRNAs and their promoters in human and mouse. Nat. Biotechnol. 35, 872–878 (2017).

64. Juzenas, S. et al. A comprehensive, cell specific microRNA catalogue of human peripheral blood. Nucleic Acids Res. 45, 9290–9301 (2017).

65. McCall, M. N. et al. Toward the human cellular microRNAome. Genome Res. (2017). doi:10.1101/gr.222067.117

66. Kitano, J., Yoshida, K. & Suzuki, Y. RNA sequencing reveals small RNAs differentially expressed between incipient Japanese threespine sticklebacks. BMC Genomics 14, 214 (2013).

67. Vaz, C. et al. Deep sequencing of small RNA facilitates tissue and sex associated microRNA discovery in zebrafish. BMC Genomics 16, 950 (2015).

68. Andreassen, R. et al. Discovery of miRNAs and Their Corresponding miRNA Genes in Atlantic Cod (Gadus morhua): Use of Stable miRNAs as Reference Genes Reveals Subgroups of miRNAs That Are Highly Expressed in Particular Organs. Plos ONE 11, e0153324 (2016).

69. Miska, E. A. et al. Microarray analysis of microRNA expression in the developing mammalian brain. Genome Biol. 5, R68 (2004).

70. Obernosterer, G., Leuschner, P. J. F., Alenius, M. & Martinez, J. Post-transcriptional regulation of microRNA expression. RNA 12, 1161–1167 (2006).

71. Makeyev, E. V., Zhang, J., Carrasco, M. A. & Maniatis, T. The MicroRNA miR-124 Promotes Neuronal Differentiation by Triggering Brain-Specific Alternative Pre-mRNA Splicing. Mol. Cell 27, 435–448 (2007).

72. Cheng, L.-C., Pastrana, E., Tavazoie, M. & Doetsch, F. miR-124 regulates adult neurogenesis in the subventricular zone stem cell niche. Nat. Neurosci. 12, 399–408 (2009).

73. Jung, H.-J. et al. Regulation of prelamin A but not lamin C by miR-9, a brain-specific microRNA. Proc. Natl. Acad. Sci. 109, E423–E431 (2012).

74. Chen, J.-F. et al. The role of microRNA-1 and microRNA-133 in skeletal muscle proliferation and differentiation. Nat. Genet. 38, 228–233 (2006).

75. Bhuiyan, S. S. et al. Evolution of the myosin heavy chain gene MYH14 and its intronic microRNA miR-499: muscle-specific miR-499 expression persists in the absence of the ancestral host gene. BMC Evol. Biol. 13, 142 (2013).

76. Horak, M., Novak, J. & Bienertova-Vasku, J. Muscle-specific microRNAs in skeletal muscle development. Dev. Biol. 410, 1–13 (2016).

77. Siddique, B. S., Kinoshita, S., Wongkarangkana, C., Asakawa, S. & Watabe, S. Evolution and Distribution of Teleost myomiRNAs: Functionally Diversified myomiRs in Teleosts. Mar. Biotechnol. 18, 436–447 (2016).

78. Cutting, A. D. et al. The potential role of microRNAs in regulating gonadal sex differentiation in the chicken embryo. Chromosome Res. 20, 201–213 (2012).

79. Muñoz, X., Mata, A., Bassas, L. & Larriba, S. Altered miRNA Signature of Developing Germ-cells in Infertile Patients Relates to the Severity of Spermatogenic Failure and Persists in Spermatozoa. Sci. Rep. 5, 17991 (2015).

80. Wainwright, E. N. et al. SOX9 Regulates MicroRNA miR-202-5p/3p Expression During Mouse Testis Differentiation. Biol. Reprod. 89, (2013).

81. Zhang, J. et al. MiR-202-5p is a novel germ plasm-specific microRNA in zebrafish. Sci. Rep. 7, 7055 (2017).

82. Gay, S. et al. MicroRNA-202 (miR-202) controls female fecundity by regulating medaka oogenesis. bioRxiv 287359 (2018). doi:10.1101/287359

83. Helfman, G., Collette, B. B., Facey, D. E. & Bowen, B. W. The Diversity of Fishes: Biology, Evolution, and Ecology, 2nd Edition. (Wiley-Blackwell, 2009).

84. Martínez, P. et al. Genetic architecture of sex determination in fish: applications to sex ratio control in aquaculture. Front. Genet. 5, (2014).

85. Spiegel, I., Salomon, D., Erne, B., Schaeren-Wiemers, N. & Peles, E. Caspr3 and Caspr4, Two Novel Members of the Caspr Family Are Expressed in the Nervous System and Interact with PDZ Domains. Mol. Cell. Neurosci. 20, 283–297 (2002).

86. Hirata Haruna et al. Cell adhesion molecule contactin-associated protein 3 is expressed in the mouse basal ganglia during early postnatal stages. J. Neurosci. Res. 94, 74–89 (2015).

87. Hirata, H., Takahashi, A., Shimoda, Y. & Koide, T. Caspr3-Deficient Mice Exhibit Low Motor Learning during the Early Phase of the Accelerated Rotarod Task. Plos ONE 11, e0147887 (2016).

88. Tong, D. et al. The critical role of ASD-related gene CNTNAP3 in regulating synaptic development and social behavior in mice. bioRxiv 260083 (2018). doi:10.1101/260083

89. Okita, M., Yoshino, Y., Iga, J. & Ueno, S. Elevated mRNA expression of CASPR3 in patients with schizophrenia. Nord. J. Psychiatry 71, 312–314 (2017).

90. Thisse, B. & Thisse, C. Fast Release Clones: A High Throughput Expression Analysis. ZFIN Direct Data Submiss. (2004).

